# Broad human and animal coronavirus neutralisation by SARS-CoV-2 S2-targeted vaccination

**DOI:** 10.1101/2021.11.30.470568

**Authors:** Kevin W. Ng, Nikhil Faulkner, Katja Finsterbusch, Mary Wu, Ruth Harvey, Saira Hussain, Maria Greco, Yafei Liu, Svend Kjaer, Charles Swanton, Sonia Gandhi, Rupert Beale, Steve J. Gamblin, Peter Cherepanov, John McCauley, Rodney Daniels, Michael Howell, Hisashi Arase, Andeas Wack, David L.V. Bauer, George Kassiotis

**Author notes:** Equal contribution (Kevin W. Ng, Nikhil Faulkner).

## Abstract

Several common-cold coronaviruses (HCoVs) are endemic in humans and several variants of severe acute respiratory syndrome coronavirus 2 (SARS-CoV-2) have emerged during the current Coronavirus disease 2019 (COVID-19) pandemic. Whilst antibody cross-reactivity with the Spike glycoproteins (S) of diverse coronaviruses has been documented, it remains unclear whether such antibody responses, typically targeting the conserved S2 subunit, contribute to or mediate protection, when induced naturally or through vaccination. Using a mouse model, we show that prior HCoV-OC43 S immunity primes neutralising antibody responses to otherwise subimmunogenic SARS-CoV-2 S exposure and promotes S2-targeting antibody responses. Moreover, mouse vaccination with SARS-CoV-2 S2 elicits antibodies that neutralise diverse animal and human alphacoronaviruses and betacoronaviruses *in vitro*, and protects against SARS-CoV-2 challenge *in vivo*. Lastly, in mice with a history of SARS-CoV-2 Wuhan-based S vaccination, further S2 vaccination induces stronger and broader neutralising antibody response than booster Wuhan S vaccination, suggesting it may prevent repertoire focusing caused by repeated homologous vaccination. The data presented here establish the protective value of an S2-targeting vaccine and support the notion that S2 vaccination may better prepare the immune system to respond to the changing nature of the S1 subunit in SARS-CoV-2 variants of concern (VOCs), as well as to unpredictable, yet inevitable future coronavirus zoonoses.

## Introduction

The ongoing Coronavirus disease 2019 (COVID-19) pandemic has highlighted the potential pathogenicity of coronaviruses (CoVs), as well as the gaps in our understanding of immune-mediated protection against this type of infection. The incidence and severity of disease varies widely following infection with acute respiratory syndrome coronavirus 2 (SARS-CoV-2), the etiologic agent of COVID-19, for reasons that remain incompletely understood *(1-3)*. In contrast, four endemic human coronaviruses (HCoVs) follow a seasonal pattern of circulation and, while they can cause significant disease in vulnerable populations, they are typically associated with common colds *(4, 5)*.

The initial successes of COVID-19 vaccination campaigns in reducing both the risk of infection with SARS-CoV-2 and the severity of symptoms, even when infection is not prevented, underscores the contribution of adaptive immunity in shaping SARS-CoV-2 pathogenicity. Indeed, in highly immune populations, SARS-CoV-2 pathogenicity may eventually fall to levels comparable with those of HCoVs *(6)*. However, the precise nature or targets of a protective immune response to SARS-CoV-2 or CoVs more broadly are only now beginning to emerge *(2, 3)*, with humoral immunity considered the mainstay *(7)*. Yet, despite almost universal seropositivity, HCoVs infect repeatedly *(8)*. Similarly, SARS-CoV-2 reinfections in previously infected individuals and breakthrough SARS-CoV-2 infections in fully COVID-19 vaccinated individuals have been documented *(9, 10)*. These observations indicate that sterilising immunity to CoVs, at least the type that is induced by natural infection or current vaccines, is partial and short-lived.

The effectiveness of vaccine-induced immunity to SARS-CoV-2 is further reduced by the emergence of virus variants that are relatively resistant to neutralisation by antibodies raised against the Spike (S) protein of the earliest SARS-CoV-2 strain isolated in Wuhan, China, which forms the basis of currently licensed vaccines *(11-19)*. Both the SARS-CoV-2 strain carrying the S D614G substitution that quickly replaced the Wuhan strain early during the pandemic and the Alpha strain (B.1.1.7) that arose in the United Kingdom exhibit modest resistance to neutralisation by antibodies raised against the Wuhan strain. Several-fold more resistant, however, are the current SARS-CoV-2 variants of concern (VOCs), Beta (B.1.351), Gamma (B.1.1.28) and Delta (B.1.617.2), first detected in South Africa, Brazil and India, respectively *(11-19)*.

Evasion of immunity by VOCs is attributable to amino acid substitutions predominantly in the S1 subunit of the S protein, a subunit that is far less conserved than the S2 subunit. Of all the substitutions in the S proteins of the Alpha, Beta, Gamma and Delta variants, 29 are located in S1, 6 of which are shared among at least two of the variants, and 6 are located in S2, none of which are shared.

As part of full-length S, the S2 subunit harbours a sizeable proportion of epitopes targeted by antibodies in the response to SARS-CoV-2 infection or vaccination *(20-27)*. However, despite the higher sequence conservation in S2 among SARS-CoV-2 variants and HCoVs, as well as all animal CoVs that have been studied, the antibody repertoire appears to focus on more variable and likely more immunogenic epitopes in the receptor binding domain (RBD) and N-terminal domain (NTD) of S1 upon repeated infections with homologous and heterologous HCoVs or SARS-CoV-2 strains, and this can vary widely between individuals *(26, 27)*. Nevertheless, cross-reactive antibodies targeting conserved regions in SARS-CoV-2 S2, such as the fusion peptide, heptad repeats (HRs), stem helix or membrane proximal regions have been detected in pre-pandemic sera *(28-31)*, likely induced by HCoVs, and are also back-boosted by SARS-CoV-2 infection or vaccination *(27, 29, 31-33)*.

As S2-targeting antibodies do not directly affect binding of the RBD to the main SARS-CoV-2 cellular receptor angiotensin converting enzyme 2 (ACE2), their relative contribution to protection and, by extension, the limits of cross-reactive immunity, have been a matter of debate. However, several S2-specific monoclonal antibodies with potent neutralising activity have been isolated during the previous SARS-CoV pandemic *(34-36)*, and more recently during the current SARS-CoV-2 pandemic *(37-42)*, implying that broad protection against diverse CoVs may be possible by targeting the S2 subunit *(43)*.

Here, we used a mouse model to directly test the contribution of S2-targeting humoral immunity to protection against SARS-CoV-2. We show that prior HCoV-OC43 S immunisation primes neutralising antibody responses to a single SARS-CoV-2 S immunisation, which would otherwise be subimmunogenic, and focuses the antibody response to S2. We further show that S2-based vaccination induces robust *in vitro* neutralisation of all human and animal CoVs tested, as well as efficient *in vivo* protection against SARS-CoV-2 challenge. Thus, S2 vaccination broadens the antibody response to highly conserved epitopes, forming the basis of a potential pan-CoV vaccine.

## Results

### Prior HCoV-OC43 S immunity cross-reacts with and primes SARS-CoV-2 S antibody responses

To examine the extent of cross-reactivity between a HCoV S and SARS-CoV-2 S, we immunised C57BL/6J mice with HCoV-OC43 S, using two doses of a DNA vaccine administered four weeks apart (**Fig. 1A**; Methods). A single dose of the vaccine (prime) did not induce SARS-CoV-2 RBD or S1 cross-reactive antibodies and only low levels of SARS-CoV-2 S2 cross-reactive antibodies, as detected by ELISA (**Fig. 1B**). The second dose of the vaccine (boost) increased the levels of these antibodies, particularly for SARS-CoV-2 S2, which were boosted 2.5-times (**Fig. 1B**). HCoV-OC43 S immunisation also induced neutralising activity against retroviral vectors pseudotyped with SARS-CoV-2 S, which was, however, one order of magnitude lower than that against homologous HCoV-OC43 S pseudotypes (**Fig. 1C**). It also induced weak neutralising activity against authentic SARS-CoV-2 Wuhan and Alpha strains, which was detected by a plaque reduction neutralisation test (PRNT) only in some of the mice and only at the highest serum concentration (**Fig. S1**).

**Figure 1.**
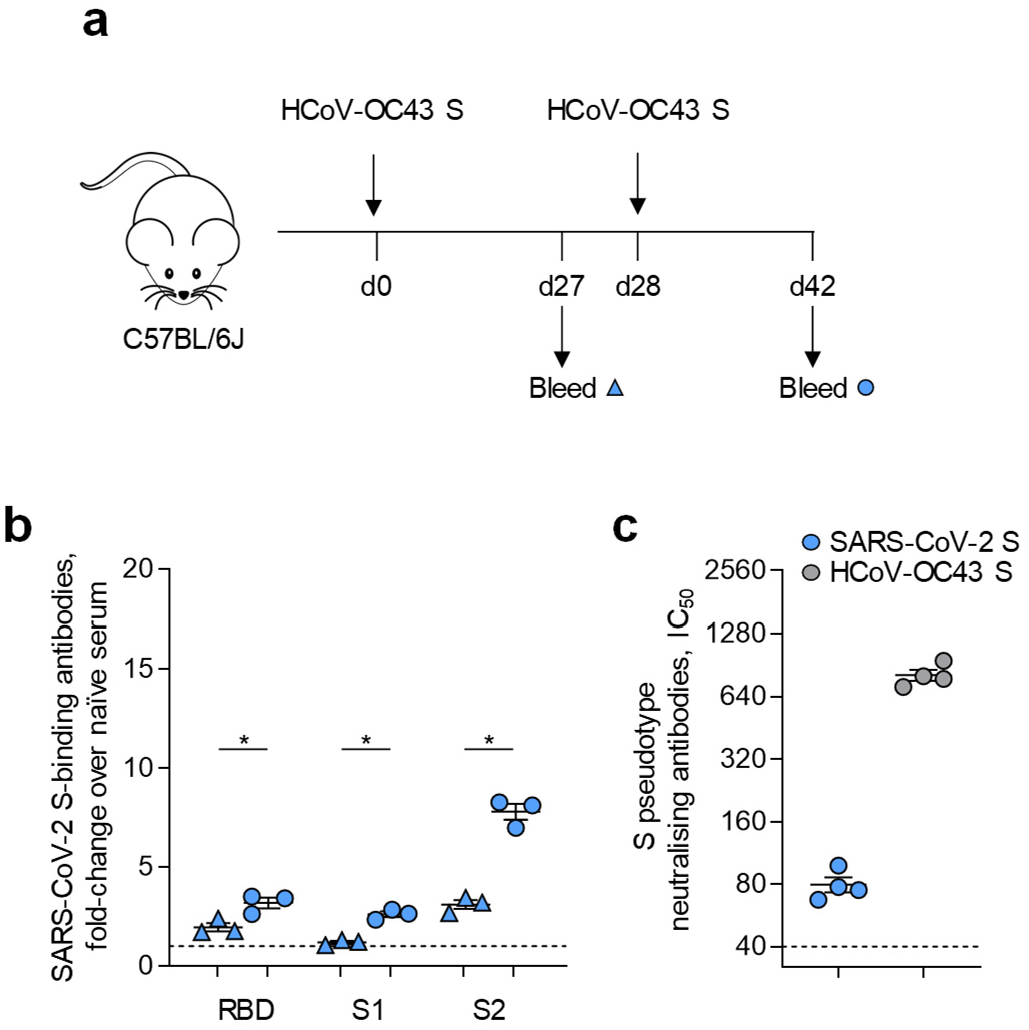
Prior HCoV-OC43 S immunity cross-reacts with SARS-CoV-2 S. **A**, Diagram of HCoV-OC43 S immunisation regimen and serum sample collection. **B**, Levels of ELISA detected antibodies reacting with SARS-CoV-2 RBD, S1 or S2 in C57BL/6J mice (n=3) after one or two doses of a HCoV-OC43 S DNA vaccine. P values were calculated with paired Student’s t-tests. **C**, Levels (IC_50_) of antibodies in the sera of mice after two doses of a HCoV-OC43 S DNA vaccine (n=4), compared with sera from unvaccinated mice (n=4), able to neutralise retroviral vectors pseudotyped with HCoV-OC43 S or SARS-CoV-2 S. Values in B and C are from two different experiments.

Another measure of S cross-reactivity is the ability of one S variant to prime responses to another. To test this, we compared the responses to a single SARS-CoV-2 Wuhan S immunisation of C57BL/6J mice that were previously immunised once with HCoV-OC43 S (prime) or not (**Fig. 2A**). As controls, we used mice immunised twice with either HCoV-OC43 S or SARS-CoV-2 Wuhan S (prime – boost) (**Fig. 2A**). Again, compared with naïve mice, a single dose of HCoV-OC43 S induced very low levels of SARS-CoV-2 S cross-reactive antibodies (**Fig. 2B**). As expected, a single dose of SARS-CoV-2 Wuhan S given to previously naïve mice induced ELISA detectable antibodies that reacted with homologous SARS-CoV-2 RBD, S1 and S2 at comparable levels (**Fig. 2B**). However, in HCoV-OC43 S primed mice, a single dose of SARS-CoV-2 Wuhan S induced significantly higher levels specifically of SARS-CoV-2 S2-binding antibodies, which were boosted 1.9-times (**Fig. 2B**). Prior HCoV-OC43 S immunisation significantly increased the levels of IgG antibodies elicited by a single dose of a subsequent SARS-CoV-2 Wuhan S vaccine that bound natural SARS-CoV-2 S conformations on a cell-based assay (**Fig. 2C**). Moreover, prior HCoV-OC43 S immunisation primed production of neutralising antibodies against authentic SARS-CoV-2 Wuhan and D614G strains, as well as the VOCs tested, detected by a high-throughput WHO-benchmarked neutralisation assay (**Fig. 2D**). In stark contrast, a single dose of the SARS-CoV-2 Wuhan S vaccine induced no measureable neutralising antibodies against any of the viruses in previously naïve mice (**Fig. 2D**). Together, these results demonstrated that prior HCoV-OC43 S exposure imprinted preferential targeting of the S2 subunit during subsequent SARS-CoV-2 S exposure and enabled the induction of neutralising antibodies that would not have otherwise been induced by a single SARS-CoV-2 S vaccine dose, thus transforming subimmunogenic SARS-CoV-2 S exposure to immunogenic.

**Figure 2.**
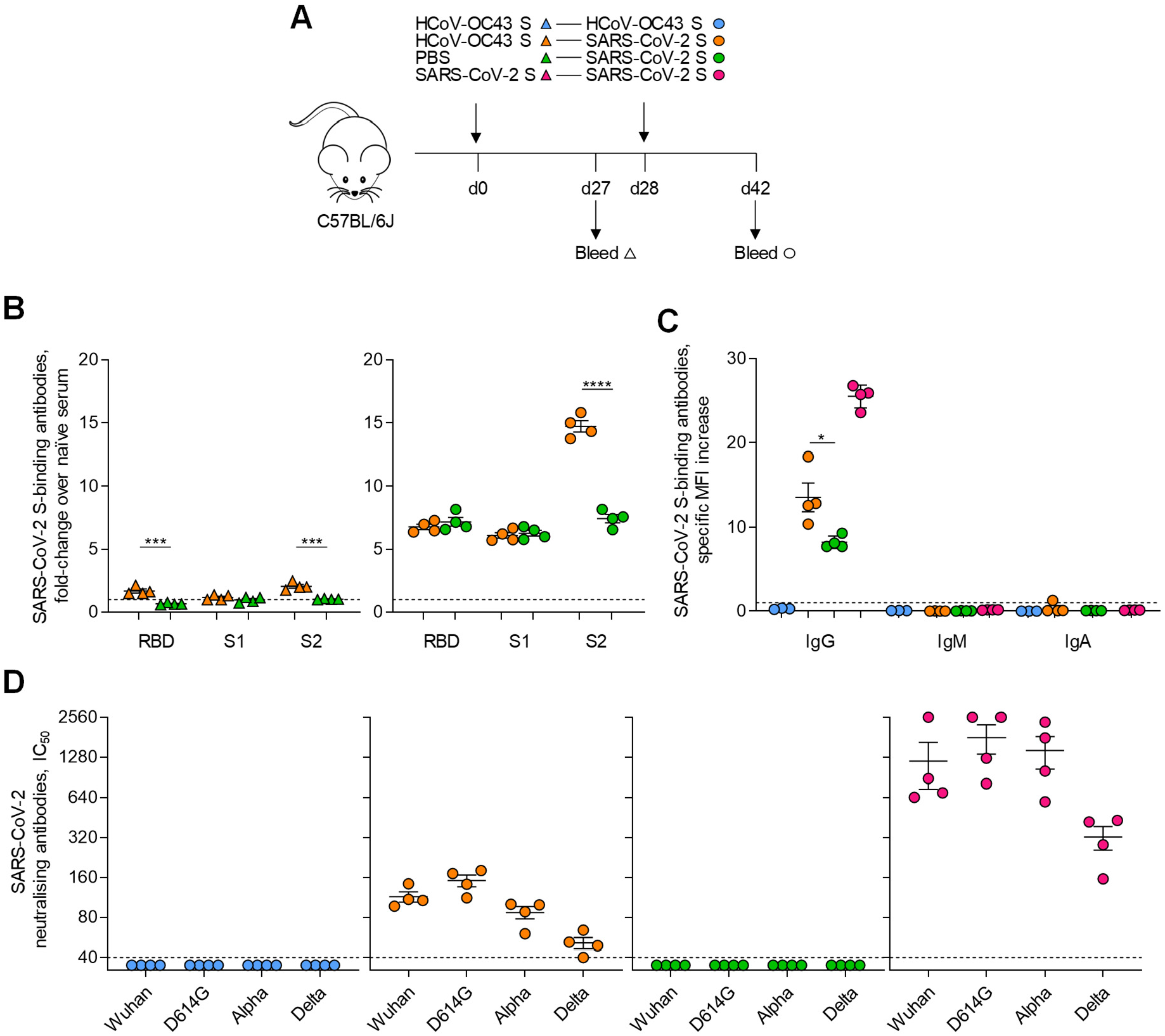
Prior HCoV-OC43 S immunity primes antibody responses to SARS-CoV-2 S. **A**, Diagram of HCoV-OC43 S and SARS-CoV-2 immunisation regimens and serum sample collection. **B**, Levels of ELISA detected antibodies reacting with SARS-CoV-2 RBD, S1 or S2 in naïve or HCoV-OC43 S-primed C57BL/6J mice (left) and after boost with SARS-CoV-2 S (right) (n=4 per group). **C**, Levels of IgG, IgM and IgA antibodies reacting with SARS-CoV-2 S in the SUP-T1 cell-based assay, in the sera of mice two weeks after the indicated prime-boost regimen (n=4 for all groups). **D**, Levels (IC_50_) of antibodies in the same groups of mice able to neutralise the indicated authentic SARS-CoV-2 strains and VOCs (n=4 per group).

### Immunisation with membrane-bound SARS-CoV-2 S2 induces broadly neutralising antibodies

As HCoV-OC43 S priming enhanced the production both of S2-targeting and of neutralising antibodies by subsequent SARS-CoV-2 S immunisation, we next explored if the two were linked. Levels of S2-targeting antibodies could simply reflect levels of RBD targeting neutralising antibodies or they could mediate at least part of the increased neutralising activity. To directly test the ability of S2-targeting antibodies to neutralise SARS-CoV-2 and other CoVs, we immunised C57BL/6J mice with a DNA vaccine encoding the membrane-bound SARS-CoV-2 S2 subunit *(20)* or with recombinant monomeric SARS-CoV-2 S2 protein. Compared with a DNA vaccine encoding the full-length SARS-CoV-2 S, both the S2 DNA vaccine and the recombinant S2 protein induced over 4-fold higher levels of S2-targeting antibodies as detected by S2-specific ELISA (**Fig. 3A**), indicating that full-length S may stimulate only a fraction of the potential magnitude of the response to S2. However, pertinent to potential neutralisation, only antibodies induced by the S2 DNA vaccine, but not by the recombinant S2 protein, recognised the intact SARS-CoV-2 S in its natural conformation in the cell-based assay (**Fig. 3B**), suggesting that the S2 construct expressed by the DNA vaccine retained naturally presented epitopes in the full-length S.

**Figure 3.**
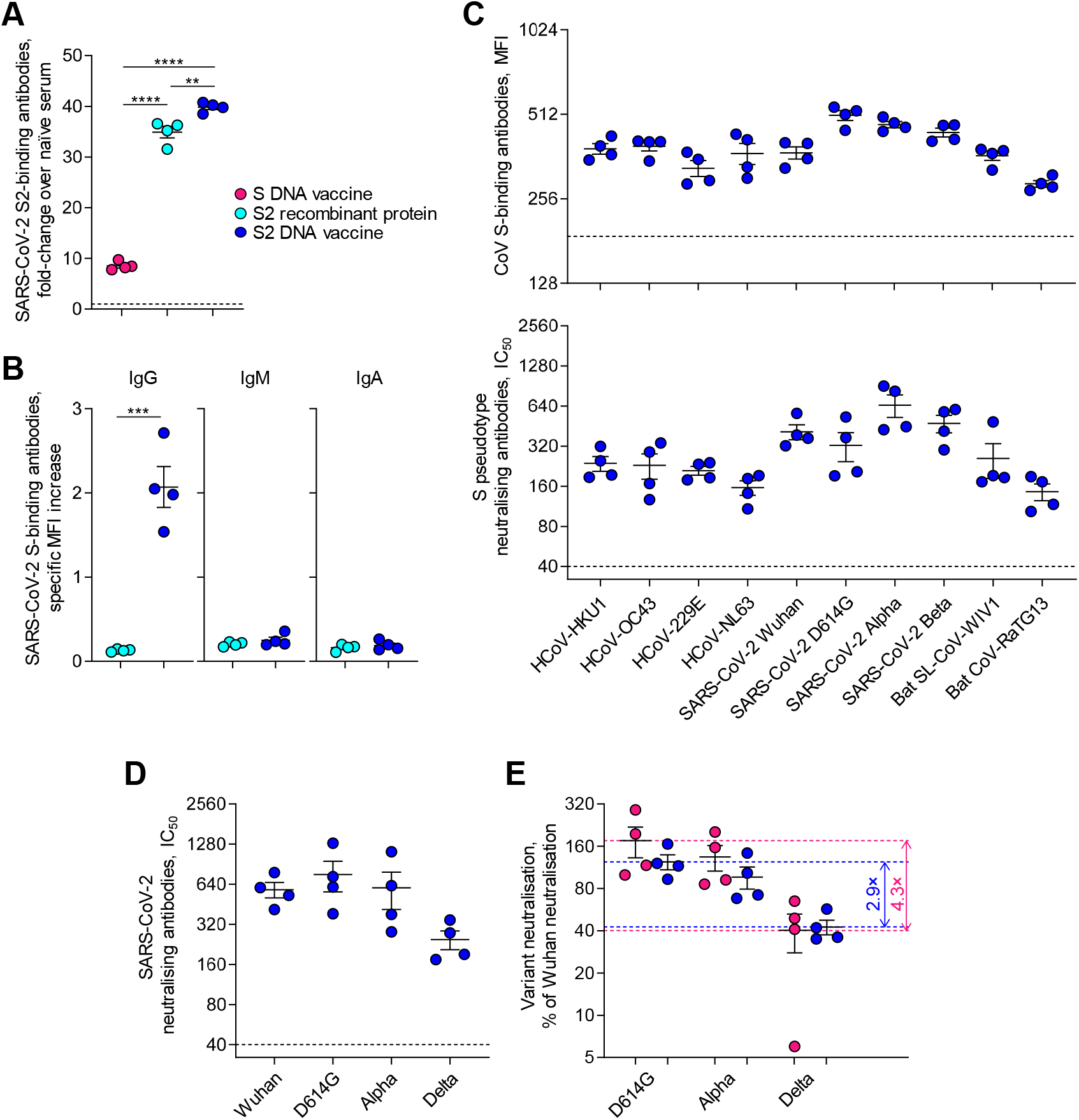
Immunisation with membrane-bound SARS-CoV-2 S2 induces broadly neutralising antibodies. **A**, Levels of ELISA detected antibodies reacting with SARS-CoV-2 S2 in the sera of C57BL/6J mice (n=4 per group) after two immunisations with DNA vaccines encoding SARS-CoV-2 full-length S or S2 or with recombinant S2 protein. **B**, Levels of IgG, IgM and IgA antibodies reacting with SARS-CoV-2 full-length S in the SUP-T1 cell-based assay in the sera of mice (n=4 per group) after two immunisations with a SARS-CoV-2 S2 DNA vaccine or with recombinant S2 protein. **C**, Levels of antibodies in the sera of mice (n=4 per group) after two immunisations with a SARS-CoV-2 S2 DNA vaccine that react with the S proteins of the indicated human and animal CoVs in the HEK293T cell-based assay (top) and corresponding neutralisation titres with retroviral vectors pseudotyped with the same S proteins (bottom). **D**, Levels (IC_50_) of antibodies in the same mice as in C, able to neutralise the indicated authentic SARS-CoV-2 strains and VOCs. **E**, Neutralisation activity against heterologous SARS-CoV-2 strains and VOCs, expressed as a percentage of neutralisation of the homologous Wuhan strain, in the sera of mice that received two immunisations with DNA vaccines encoding SARS-CoV-2 full-length S or S2 (n=4 per group). Data from a single experiment are shown. SARS-CoV-2 S2 immunisation was repeated once more with similar results. Horizontal dashed lines mark the average neutralising activity for the maximally and minimally neutralised group and numbers represent the ratio between the two.

Reactivity of SARS-CoV-2 S2-immune mouse sera with short overlapping peptides in an array covering the SARS-CoV-2 S2 subunit revealed a similar pattern to that with COVID-19 convalescent human sera *(28)*, but the mouse sera reacted less well with epitopes in the fusion peptide region than with other defined epitopes in S2 (**Fig. S2**).

Antibodies induced by SARS-CoV-2 S2 immunisation recognised the full-length S not only from the homologous SARS-CoV-2 Wuhan strain but also from all 4 HCoVs, other SARS-CoV-2 strains and VOCs, and the two bat CoVs, bat SARS-like coronavirus WIV1 (Bat SL-CoV-WIV1) and bat coronavirus RaTG13 (Bat CoV-RaTG13) as tested by flow cytometry in a cell-based assay (**Fig. 3C**). Furthermore, these antibodies effectively neutralised retroviral vectors pseudotyped with all the CoV S proteins tested (**Fig. 3C**), indicating broad neutralising ability. In the high-throughput authentic virus neutralisation assay, antibodies induced by SARS-CoV-2 S2 immunisation demonstrated neutralising activity against SARS-CoV-2 Wuhan and D614G strains, as well as the VOCs tested (**Fig. 3D**), at levels comparable with those induced by full-length S immunisation (**Fig. 2D**). Relative to recognition of the homologous Wuhan strain, S2 immunisation induced neutralising antibodies that were less sensitive to amino acid substitutions in SARS-CoV-2 S variants than full-length S immunisation, as suggested by the difference between the maximally and minimally neutralised SARS-CoV-2 variant, which we consider as an indirect measure of cross-reactivity (**Fig. 3C**). Therefore, immunisation with cell-presented membrane-bound SARS-CoV-2 S2, but not with recombinant S2 protein, induced neutralising antibodies with broad activity against all animal and human CoVs tested.

### Immunisation with membrane-bound SARS-CoV-2 S2 affords *in vivo* protection

Titres of neutralising antibodies elicited by SARS-CoV-2 S2 immunisation were determined also with an authentic virus high-throughput neutralisation assay *(44)*, which been used to establish the WHO International Standard for SARS-CoV-2 antibody neutralisation and was benchmarked against this standard *(19)*. The half-maximal inhibitory concentration (IC_50_) in this benchmarked assay *(18, 19)* and in similar assays has been correlated with vaccine efficacy against SARS-CoV-2, including the Delta variant in humans *(45-47)*, suggesting that the neutralising antibody titres that SARS-CoV-2 S2 immunisation induced in mice would also be protective against viral infection *in vivo*. Nevertheless, to directly test the level of *in vivo* protection afforded by SARS-CoV-2 S2 immunisation, we next challenged SARS-CoV-2 S2 vaccinated and control unvaccinated mice with SARS-CoV-2 Wuhan or Alpha variants. Although certain SARS-CoV-2 variants, particularly ones bearing the S N501Y substitution, can replicate in wild-type C57BL/6J mice using the murine ACE2 as cellular receptor, for this study, we employed K18-hACE2 transgenic mice expressing human ACE2 in epithelial cells, under the control of the human keratin 18 promoter *(48)*, which may better capture the ability of antibodies to disrupt virus replication using the human ACE2 receptor. Given the severe pathology that results from SARS-CoV-2 infection of K18-hACE2 transgenic mice at later time-points, infected mice were analysed 4 days after challenge by expression of the SARS-CoV-2 envelope (E) gene in infected lungs (**Fig. 4A**). In contrast to unvaccinated controls, which had readily detectable levels of SARS-CoV-2 E RNA in their lungs, SARS-CoV-2 S2 vaccinated mice were strongly protected against both SARS-CoV-2 Wuhan and Alpha challenge (**Fig. 4B**), supporting a protective effect of SARS-CoV-2 S2 immunisation, as inferred from *in vitro* neutralisation data.

**Figure 4.**
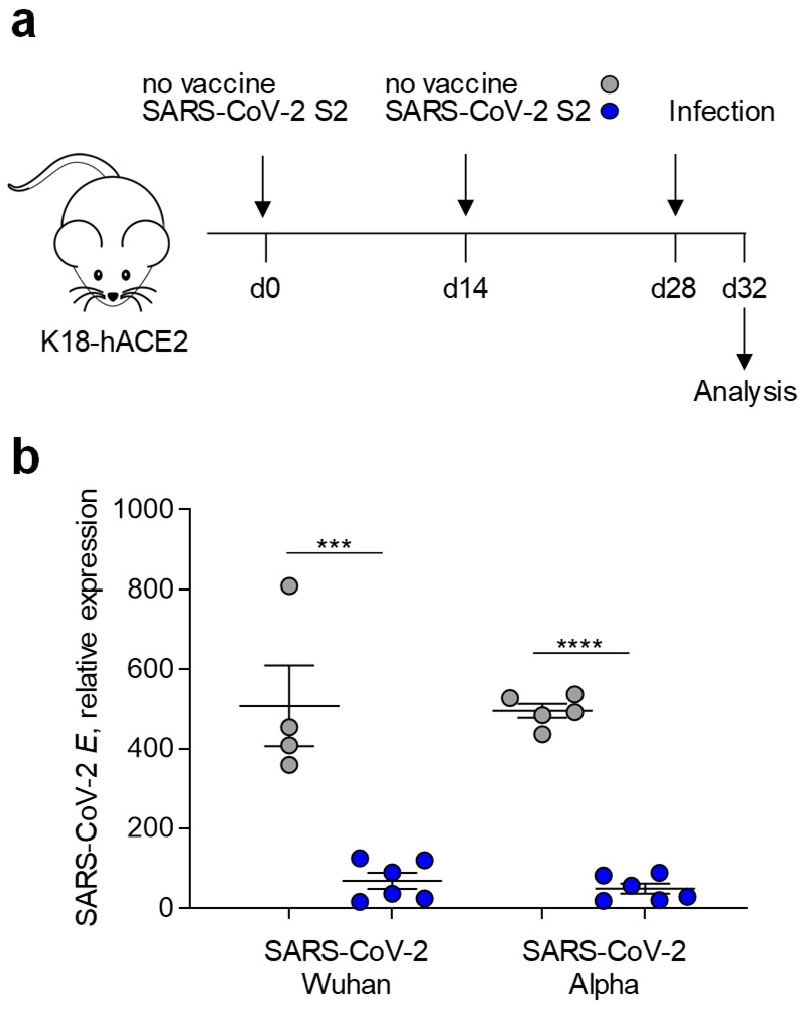
Immunisation with membrane-bound SARS-CoV-2 S2 protects against *in vivo* challenge. **A**, Diagram of SARS-CoV-2 S2 immunisation regimen and SARS-CoV-2 challenge. **B**, Virus loads, determined by RT-qPCR for SARS-CoV-2 E expression, in whole lungs from K18-hACE2 transgenic mice that were either unvaccinated (n=4-5) or vaccinated with SARS-CoV-2 S2 (n=6) and were subsequently challenged with SARS-CoV-2 Wuhan or Alpha.

### SARS-CoV-2 S2 immunisation boosts prior SARS-CoV-2 S-induced cross-reactivity

Repeated immunisation with homologous antigens, such as the Influenza A vaccine, may boost antibody responses to immunodominant, but serotype-specific epitopes, at the expense of less dominant but conserved epitopes, thus progressively narrowing the antibody repertoire and cross-reactivity with heterologous strains *(49)*. In standard COVID-19 vaccination campaigns, individuals receive two doses of a SARS-CoV-2 S Wuhan-based vaccine and certain groups now receive a third dose of the same or similar vaccines, carrying the same antigen. Although repeated homologous vaccination boosts overall antibody titres, at least transiently, its effect on cross-reactivity with heterologous strains has not yet been established. To mimic at least part of the immunological history imprinted by current vaccination campaigns, we administered two doses of the SARS-CoV-2 S Wuhan vaccine before examining the response to a third dose of either the same or an alternative S2-targeted vaccine.

A group of C57BL/6J mice were immunised twice with the SARS-CoV-2 S Wuhan DNA vaccine two weeks apart and were then split into two groups, each of which received either a third dose of the same vaccine or the SARS-CoV-2 S2 vaccine (**Fig. 5A**). Two weeks after the third immunisation, all sera demonstrated strong neutralisation of the immunising Wuhan strain, as well as the D614G strain and Alpha VOC, which was significantly higher after the S2 vaccine than after the third dose of the full-length S vaccine (**Fig. 5B**). Differences were more pronounced in neutralisation of the Delta VOC, which was generally more resistant to neutralisation, with 2 out of 6 mice in the full-length S vaccine arm lacking detectable levels of Delta neutralising activity (**Fig. 5B**). Even when neutralising antibody titres were normalised to the neutralisation of the homologous Wuhan strains, S2 booster immunisation elicited more uniform activity against the VOCs tested than full-length S booster immunisation, again indicating broader cross-reactivity of the former response (**Fig. 5C**). Together, these results suggested that S2 immunisation induces robust immunity against SARS-CoV-2 VOCs even against the backdrop of repeated SARS-CoV-2 full-length Wuhan S immunisation.

**Figure 5.**
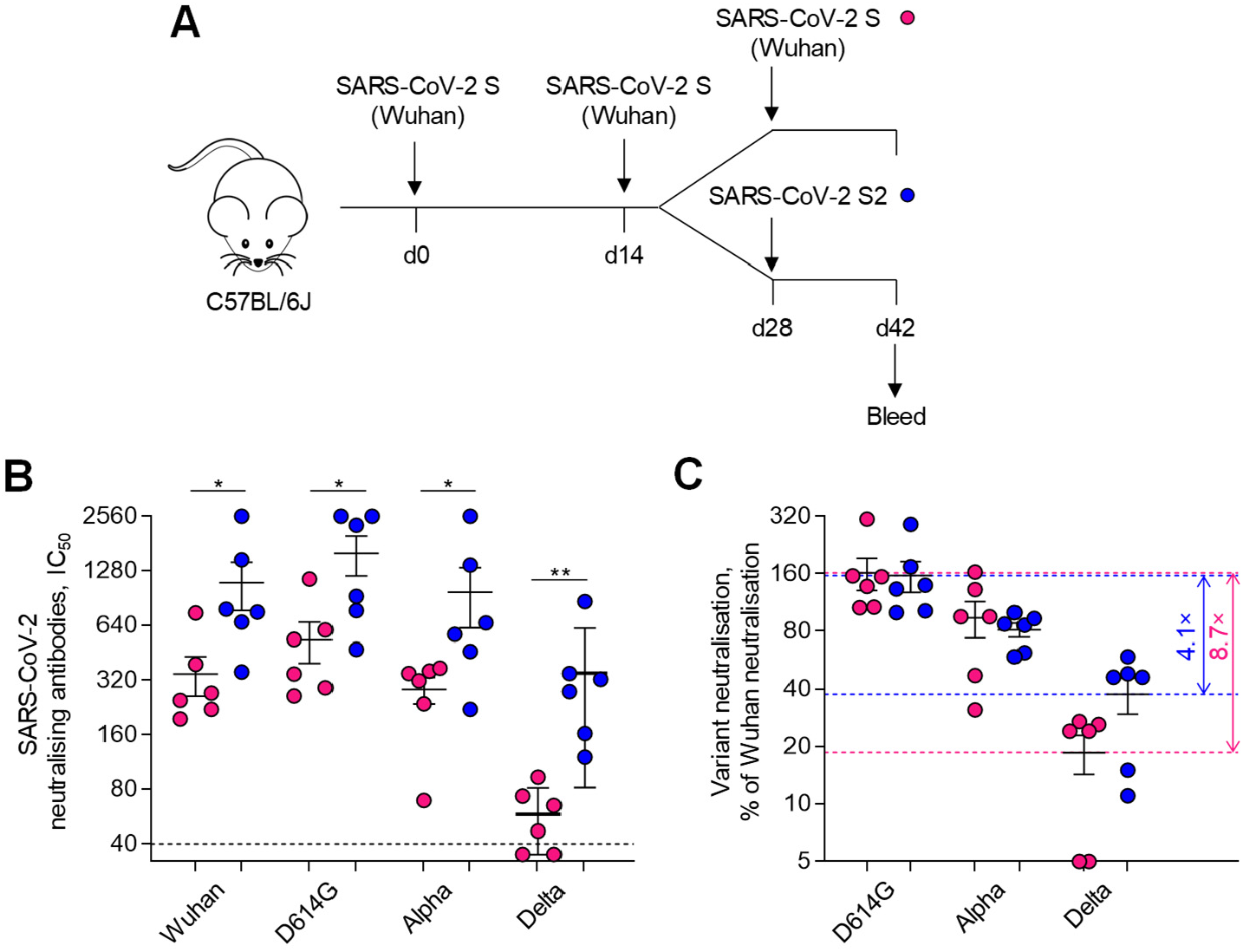
SARS-CoV-2 S2 immunisation boosts prior SARS-CoV-2 S-induced cross-reactivity. **A**, Diagram of SARS-CoV-2 full-length S and S2 immunisation regimens and serum sample collection. **B**, Levels (IC_50_) of antibodies able to neutralise the indicated authentic SARS-CoV-2 strains and VOCs in the sera of mice that were immunised twice with SARS-CoV-2 full-length S and then received a third immunisation with either SARS-CoV-2 full-length S or S2 (n=6 per group). **C**, Neutralisation activity against heterologous SARS-CoV-2 strains and VOCs, expressed as a percentage of neutralisation of the homologous Wuhan strain, in the sera of the same mice. Horizontal dashed lines mark the average neutralising activity for the maximally and minimally neutralised group and numbers represent the ratio between the two.

## Discussion

Mutations acquired by SARS-CoV-2 VOCs that encode certain amino acid substitutions in S reduce the neutralising ability of antibodies elicited by natural infection with earlier strains, such as the Wuhan and D614G strains and the Alpha variant *(11-17, 44, 50-53)*, as well as of antibodies induced by current COVID-19 vaccines based on the SARS-CoV-2 Wuhan S sequences *(11-19)*. The evolution of SARS-CoV-2 S immune evasion warrants consideration of less variable or mutable targets in the S protein and of immunisation strategies eliciting broader neutralising activity. Here, we used a mouse model to test immunity afforded by targeting conserved regions of the S protein and provide evidence for a highly protective antibody response targeting the S2 subunit. This was evidenced both by prior exposure to a HCoV, which induced SARS-CoV-2 S2 cross-reactive antibodies, and by deliberate immunisation with a SARS-CoV-2 S2-encoding DNA vaccine.

Although induction of S2-targeting cross-reactive antibodies by HCoV or SARS-CoV-2 infection or COVID-19 vaccination is now well-recognised *(27-33, 37, 39-42)*, their potential contribution to the outcome or severity of SARS-CoV-2 infection remains to be established *(43)*. A potential protective effect mediated by such antibodies is suggested by certain studies linking an early S2-targeted antibody response to SARS-CoV-2 with milder symptoms and survival of COVID-19 *(54-59)*. However, detrimental effects of cross-reactive or pre-existing immunity, such as original antigenic sin, have also been hypothesised, although not always directly attributed to S2-targeting antibodies *(20, 60-62)*.

Our observation that prior exposure of immunologically naïve mice to HCoV-OC43 S promotes S2-targeting cross-reactive antibodies and primes for induction of SARS-CoV-2 neutralising antibodies that would not have otherwise been elicited after a single SARS-CoV-2 immunisation provides direct evidence for a cross-protective effect. No evidence consistent with original antigenic sin was obtained in this model. In contrast, Lapp et al. observed that priming of mice with HCoV-HKU1 induced SARS-CoV-2 S cross-reactive antibodies, but impeded the neutralising antibody response to subsequent SARS-CoV-2 immunisation *(63)*. However, in contrast to the mouse experiment, they also found that pre-existing HCoV-HKU1 antibodies correlated positively with SARS-CoV-2 neutralising antibodies in children after natural infection *(63)*. While it is theoretically possible that different HCoVs either impede or promote a subsequent antibody response to SARS-CoV-2, an alternative explanation resides in the fact that Lapp et al. used recombinant proteins for immunisation of mice *(63)*. In contrast to immunisation with RNA or DNA vectors or natural infection, both of which produce the natural S conformations presented on the surface of cells or virions, recombinant protein immunisation may not correctly present the targets of neutralising antibodies. Indeed, immunisation with recombinant SARS-CoV-2 S2 in this study induced high titres of antibodies able to bind recombinant S2 on ELISA, but unable to bind the full-length S presented on cells. In support of this notion, nearly one in three antibodies cloned from a SARS-CoV convalescent donor that cross-reacted with SARS-CoV-2 S, recognised the full-length S expressed on the surface of cells, but not recombinant S ectodomain *(64)*.

Contrasting with recombinant S2 protein, the SARS-CoV-2 S2-encoding vaccine used here induced a highly protective and broadly neutralising response. Eliciting protective antibodies against the S2 subunit has been attempted already during the SARS-CoV epidemic in 2005, but not always successfully. Guo et al. reported that immunisation of mice with either a recombinant SARS-CoV S2 fragment or a DNA vaccine encoding SARS-CoV S2 induced high titres of S2-binding antibodies, but no SARS-CoV neutralising activity *(65)*. Using a smaller recombinant SARS-CoV S2 fragment, encompassing the connector domain, stem helix, HR2 and membrane proximal regions (corresponding to S_1073-1210_ of SARS-CoV-2 S), Keng et al. elicited strong neutralising activity against SARS-CoV in immunised rabbits *(66)*, and isolated neutralising monoclonal antibodies targeting separate epitopes in the stem helix, HR2 and membrane proximal regions in a follow-up study *(34)*.

More recently, Ma et al. reported that mice immunised with a combination of nanoparticle vaccines displaying either SARS-CoV-2 RBD or an SARS-CoV-2 S2 fragment from HR1 to HR2 (S_910-1213_), but not those immunised with SARS-CoV-2 RBD alone, elicited neutralising antibodies against SARS-CoV, MERS-CoV, HCoV-229E, HCoV-OC43 and Bat CoV-RaTG13 S pseudotypes *(67)*. Although nanoparticles displaying the SARS-CoV-2 S2 fragment were not tested in isolation in this study, the findings do suggest that the observed broad neutralisation was likely mediated by S2-targeting antibodies *(67)*.

The isolation of highly neutralising S2 cross-reactive monoclonal antibodies demonstrates the potential for broad and effective targeting of conserved epitopes on S2. Several monoclonal antibodies targeting the stem helix or base of S2 have now been reported to neutralise several betacoronaviruses, but not human alphacoronaviruses, where this region of S2 is less conserved *(37, 39-41, 68)*. An antibody targeting the more conserved hinge region (S_980-1006_) of S2, however, is reported to neutralise both alphacoronaviruses and betacoronaviruses *(42)*. Despite the presence of cross-reactive epitopes, no monoclonal antibodies reactive with the fusion peptide region have been isolated to date. Nevertheless, vaccination of pigs with SARS-CoV-2 fusion peptide expressed on the surface of bacteria, induced protective immunity against the porcine epidemic diarrhea virus (PEDV), an animal CoV sharing sequence similarity in its S2 subunit with that of SARS-CoV-2 and all other CoVs *(69)*.

It is interesting to note that, to the extent indicated by the use of short overlapping peptides here, sera from S2-immunised mice appear to target the hinge region and central helix more strongly than the fusion peptide or the stem helix. These regions are considerably more conserved among human and animal CoVs than other regions of S2, and S2 as a whole is far more conserved than S1. While amino acid substitutions in S2 are rarer than in S1, they may still have significance *(23)*. For example, the D796H substitution in the highly conserved S_796_ position of S2 upstream of the fusion peptide, has been selected by immune serum therapy of an immunodeficient COVID-19 patient for its ability to evade serum neutralisation *(70)*, and it is also found in the B.1.1.318 lineage *(71)*. Resistance to neutralisation and viral rebound conferred by the D796H substitution is consistent with a significant role for S2-targeting antibodies in the control of SARS-CoV-2, but it also incurs substantial fitness cost *(70)*. Therefore, although possible, evasion of humoral immunity may be much more difficult to achieve by amino acid substitutions in critical regions of S2 without loss of virus fitness.

Independently of the precise epitopes targeted in these mice, our data provide clear evidence that S2-based immunisation can induce broad *in vitro* neutralisation of diverse alphacoronaviruses and betacoronaviruses, including human and animal CoVs. S2 immunisation also renders mice resistant to *in vivo* challenge with SARS-CoV-2, linking *in vitro* neutralisation and *in vivo* protection, but the latter may also utilise Fc-dependent antibody functions, which may contribute substantially to *in vivo* protection, particularly by S2-targeting cross-reactive antibodies *(59)*.

The data presented here argue that S2 targeting may be an important component of a pan-CoV vaccine. While it is unlikely that any CoV vaccine will induce full, life-long immunity, our findings suggest that if the antibody repertoire is primed or periodically reset to focus on more conserved regions in S2, then repeated natural exposure to endemic HCoVs can be leveraged to maintain cross-reactive responses to current or future pandemic-causing CoVs.

## Materials and Methods

### Study Design

This study was designed to assess the therapeutic potential of targeted coronavirus S2 vaccination against SARS-CoV-2 and other human and animal CoVs. We tested nucleic acid and protein-based vaccine candidates targeting SARS-CoV-2 S and its constituent subunits using a prime-boost system in wild-type or K18-hACE2 transgenic mice, and tested *in vitro* activity of serum antibodies and *in vivo* protection following SARS-CoV-2 challenge. Sample sizes for all animal experiments were estimated based on prior experience to provide statistical power whilst minimizing animal use and were not altered over the course of the study. Time-points were defined based on prior experience and were not altered over the course of the study, and mice were randomly assigned to treatment groups before the start of each experiment. Antibody binding against CoV S proteins and their subunits was assessed by ELISA against SARS-CoV-2 S subunits and flow cytometry on cells transfected with expression vectors encoding S from HCoVs, SARS-CoV-2 (including variants), and bat CoVs. *In vitro* neutralisation titres were determined using GFP-expressing retroviruses pseudotyped with various CoV S proteins, as well as two separate neutralisation assays using authentic SARS-CoV-2 viruses. For all *in vitro* analyses, at least two technical replicates were performed per biological replicate and the data represents the average of the technical replicates. Investigators performing live virus neutralisation and *in vivo* infection were blinded until after the experiment and analysis were completed. No samples were excluded from analysis.

### Mice

Wild-type C57BL/6J (WT) and K18-hACE2 transgenic mice *(48)* on a C57BL/6J background were obtained from The Jackson Laboratory, and bred and maintained at the Francis Crick Institute’s Biological Research Facility under specific pathogen-free conditions. All experiments were approved by the Francis Crick Institute’s ethical committee and conducted according to local guidelines and UK Home Office regulations under the Animals Scientific Procedures Act 1986 (ASPA; project licence numbers PCD77C6D0 and P9C468066).

### Virus isolates

SARS-CoV-2 virus isolates were obtained and propagated as previously described *(19, 28, 44, 52)*. The SARS-CoV-2 reference isolate (Wuhan) was hCoV-19/England/02/2020 obtained from Public Health England (PHE). The D614G strain (B.1.1) was isolated from a swab from an infected healthcare worker at University College London Hospital (UCLH), obtained through the SAFER study *(72)* and carries only the D614G substitution in the S protein *(19)*. The Alpha (B.1.1.7) isolate was hCoV-19/England/204690005/2020, carrying the D614G, Δ69–70, Δ144, N501Y, A570D, P681H, T716I, S982A, and D1118H substitutions, and was obtained from PHE, through Prof. Wendy Barclay, Imperial College London, London, UK and the Genotype-to-Phenotype National Virology Consortium (G2P-UK). The Delta isolate (B.1.617.2) was MS066352H (GISAID EpiCov™ accession number EPI_ISL_1731019), carrying the T19R, K77R, G142D, Δ156-157/R158G, A222V, L452R, T478K, D614G, P681R, D950N substitutions, and was kindly provided by Prof. Wendy Barclay, Imperial College London, London, UK, through G2P-UK. All viral isolates were propagated in Vero V1 cells.

### Immunisations and infections

For protein immunisation, mice were injected intraperitoneally with 25 µg recombinant SARS-CoV-2 S2 (CV2006; LifeSensors) in monophosphoryl lipid A adjuvant (Sigma Adjuvant System, Sigma Aldrich) in a 100 µl volume. For DNA immunisation, mice were injected intravenously with 100 μl of 1.6 mg/ml plasmid DNA complexed with 1.21 mM GL67 lipid (Genzyme). Serum was collected by venepuncture or terminal bleed into serum separator tubes, centrifuged at 3,000 rpm for 5 min after 30 min clotting, then centrifuged at 12,000 rpm for 5 min, heat-inactivated at 56°C for 10 min, and stored at -20°C. For *in vivo* challenge, naïve or immunised K18-hACE2 transgenic mice were infected intranasally with 3,000 plaque-forming units (pfu) SARS-CoV-2 Wuhan or Alpha strains in 50 μl PBS under light anaesthesia (3% isoflurane) in containment level 4 facilities. Pre-infection body weights were recorded, and mice were weighed daily and monitored for clinical symptoms.

### Cell lines and plasmids

HEK293T, Vero E1, Vero E6, and SUP-T1 cells were obtained from the Francis Crick Institute’s Cell Services facility and verified as mycoplasma-free. All human cell lines were further validated by DNA fingerprinting. Cells were grown in Iscove’s Modified Dulbecco’s Medium (Sigma Aldrich) with 5% fetal bovine serum (Thermo Fisher), 2 mM L-glutamine (Thermo Fisher), 100 U/ml penicillin (Thermo Fisher), and 0.1 mg/ml streptomycin (Thermo Fisher). HEK293T cells expressing coronavirus spikes were generated by transient transfection using pcDNA3 or pCMV3-based expression plasmids and GeneJuice (EMD Millipore) 48 hours prior to use, as previously described (**Fig. S3**) *(28, 44, 73)*. The expression vector (pME18S) encoding SARS-CoV-2 S2 was described previously *(20)*. Briefly, it combines a signal sequence and transmembrane domain from SLAM and PILRα proteins, respectively *(74)* with SARS-CoV-2 S_588–1219_, which covers the last 97 amino acid residues of S1 and the whole S2 ectodomain *(20)*. Expression vectors (pcDNA3) carrying a codon-optimised gene encoding the wild-type or D614G SARS-CoV-2 S (UniProt ID: P0DTC2) were kindly provided by Massimo Pizzato, University of Trento, Italy). Expression vectors (pcDNA3) carrying codon-optimised genes encoding the Alpha or Beta SARS-CoV-2 S variants *(44)*, or Bat SL-CoV-WIV1 (UniProt ID: U5WI05) or Bat CoV-RaTG13 (UniProt ID: A0A6B9WHD3) S proteins were generate in-house. Expression vectors (pCMV3) encoding HCoV-229E S (UniProt ID: APT69883.1), HCoV-NL63 S (UniProt ID: APF29071.1), HCoV-OC43 S (UniProt ID: AVR40344.1) or HCoV-HKU1 S (UniProt ID: Q0ZME7.1) were obtained from SinoBiological. SUP-T1 cells stably expressing SARS-CoV-2 S and GFP were generated as described previously *(73)*.

### Flow cytometric detection of antibodies

Serum antibodies against coronavirus spikes were quantified as described previously *(28, 73)*. Briefly, HEK293T cells were transfected to express various S proteins 48 hours prior to use, trypsinised, and dispensed into V-bottom 96-well plates (20,000-40,000 cells/well). S^+^GFP^+^ SUP-T1 cells were mixed 1:1 with parental GFP^-^ SUP-T1 cells before dispensing into V-bottom 96-well plates (20,000-40,000 total cells/well) *(73)*. Cells were incubated with sera (diluted 1:50 in PBS) for 30 min, washed with FACS buffer (PBS, 5% BSA, 0.05% sodium azide) and stained with anti-mouse IgG (Poly4053; Biolegend), IgA (Clone 11-44-2, Southern Biotech), and IgM (Clone RMM-1; Biolegend) diluted 1:200 in FACS buffer for 30 min. Cells were washed with FACS buffer and run on a Ze5 analyser (Bio-Rad) running Bio-Rad Everest software v2.4 and analysed using FlowJo v10 (Tree Star Inc.). For SUP-T1 cells, specific MFI increase caused by immune serum (**Fig. S4**) was calculated using the following formula: (MFI of S^+^ SUP-T1 – MFI of parental SUP-T1)/MFI of parental SUP-T1.

### ELISA detection of antibodies

ELISAs were run as previously described *(28, 73)* using in-house recombinant SARS-CoV-2 S1 and RBD and commercial S2 (CV2006; Life Sensors). Following incubation with serum (diluted 1:50 in PBS), plates were washed and incubated with HRP-conjugated anti-mouse IgG (ab6728; Abcam). Plates were developed by adding 50 μl TMB substrate for 5 min with shaking (Thermo Fisher), followed by 50 μl TMB stop solution (Thermo Fisher). Optical densities (ODs) were measured at 450 nm on a Tecan microplate reader, and results were expressed as fold-change of optical densities between samples and naïve serum controls.

### Pseudotyped virus neutralisation

Lentiviral particles were pseudotyped with CoV S proteins as previously described *(28)*. Briefly, HEK293T cells were transfected using GeneJuice (EMD Millipore) with plasmids encoding CoV S proteins, SIVmac Gag-Pol, and HIV-2 backbone encoding GFP. Virus-containing supernatants were collected 48 hours post-transfection and stored at -80°C until further use. For neutralisation assays, pseudotyped viruses were incubated with serial dilutions of sera at 37°C for 30 min and added to Vero E6 cells seeded in 96-well plates (3,000 cells/well). Polybrene (4 μg/ml; Sigma Aldrich) was added to the cells, which were then spun at 1,200 rpm for 45 min. GFP^+^ cells were quantified by flow cytometry 72 hours post-transduction (**Fig. S5**), and the inverse serum dilution leading to a 50% reduction in GFP^+^ cells was taken as the neutralising titre.

### Authentic virus neutralisation

SARS-CoV-2 neutralising antibody titres were determined using PRNT and high-throughput immunofluorescence assays, as previously described *(19, 28, 44)*. For PRNT assays, triplicate cultures of Vero E6 cells were incubated with SARS-CoV-2 variants and serial dilutions of heat-inactivated sera for 3 hours, the inoculum removed, and overlaid with virus growth medium containing 1.2% Avicel (FMC BioPolymer). 24 hours later, cells were fixed in 4% paraformaldehyde and permeabilised with 0.2% Triton X-100 in PBS. Virus plaques were visualised by immunostaining with a rabbit polyclonal anti-NSP8 antibody (ABIN233792; Antibodies Online) and an HRP-conjugated anti-rabbit antibody (1706515; Bio-Rad). Plaques were quantified and IC_50_ values calculated using LabView or SigmaPlot software as previously described *(28)*. For high-throughput immunofluorescence assays, Vero E6 cells were seeded in 384-well plates and incubated with SARS-CoV-2 variants and serial dilutions of heat-inactivated sera 24 hours later. Plates were incubated for 24 hours, fixed in 4% paraformaldehyde, blocked and permeabilised with 3% BSA and 0.2% Triton X-100 in PBS. Cells were immunostained with DAPI and Alexa488-conjugated anti-N antibody (CR3009; produced in-house) and imaged using an Opera Phenix (Perkin Elmer). The ratio of infected (Alexa488^+^) to total cellular (DAPI^+^) area per well was determined using Harmony (Perkin Elmer) and IC_50_ values calculated as previously described *(44)*. Performance of this high-throughput neutralisation assay has been benchmarked across laboratories participating in the establishment and validation of the WHO International Standard for SARS-CoV-2 antibody neutralisation (WHO/BS.2020.2403) *(19)*.

### RNA isolation and RT-qPCR

Mouse lungs were harvested placed in RNAlater (Ambion) and stored at -80°C. Lungs were homogenised with a FastPrep-24 5G instrument using TallPrep Lysing Matrix M tubes (MP, Cat# 116949025) in 2 ml Trizol and RNA was isolated using the RiboPure Kit (Invitrogen, Cat# AM1924), according to the manufacturer’s instructions. 500 ng RNA was reverse-transcribed using the qPCRBIO cDNA synthesis kit according to the manufacturer’s instructions. RT-qPCR was performed on an Applied Biosystems QuantStudio 3 machine using the following TaqMan primers for *Hprt1* (Mm00446968_m1): forward: 5′-ACAGGTACGTTAATAGTTAATAGCGT-3′; reverse: 5′-ATATTGCAGCAGTACGCACACA -3′; and probe: FAM-5’ ACACTAGCCATCCTTACTGCGCTTCG 3’-TAMRA. TaqMan primers for SARS-CoV-2 *E* gene were as previously described *(75)*. SARS-CoV-2 *E* values were normalised to the housekeeping gene *Hprt1*.

### Peptide array

Antibody reactivity with a peptide array (12-mers overlapping by 10 amino acid residues) spanning the last 743 amino acids of SARS-CoV-2 S was carried out as previously described *(28)*, except sera from immunised mice were used as the primary antibody and IRDye 800CW Goat anti-mouse IgG (Licor; 1:15,000 in blocking buffer) was used as a secondary antibody.

### Statistical analysis

Data were analysed and plotted in GraphPad Prism v8 (GraphPad Software) or SigmaPlot v14.0 (Systat Software). Parametric comparisons of normally-distributed values that satisfied variance criteria were made using paired or unpaired Student’s t tests or one-way ANOVA tests. Data that did not pass the variance criteria were compared using non-parametric two-tailed Mann-Whitney Rank Sum tests or ANOVA on Ranks tests.

## Acknowledgements

We are grateful for assistance from the Biological Services, Flow Cytometry and Cell Services facilities at the Francis Crick Institute and to Mr Michael Bennet and Mr Simon Caidan for training and support in the high-containment laboratory. We wish to thank the Public Health England (PHE) Virology Consortium and PHE field staff, the ATACCC (Assessment of Transmission and Contagiousness of COVID-19 in Contacts) investigators, the G2P-UK (Genotype to Phenotype-UK) National Virology Consortium, and Prof Wendy Barclay, Imperial College London, London, UK, for the B.1.1.7 virus isolate. This work was supported by the Francis Crick Institute, which receives its core funding from Cancer Research UK, the UK Medical Research Council, and the Wellcome Trust.

## Supplementary Materials for

**Figure S1.**
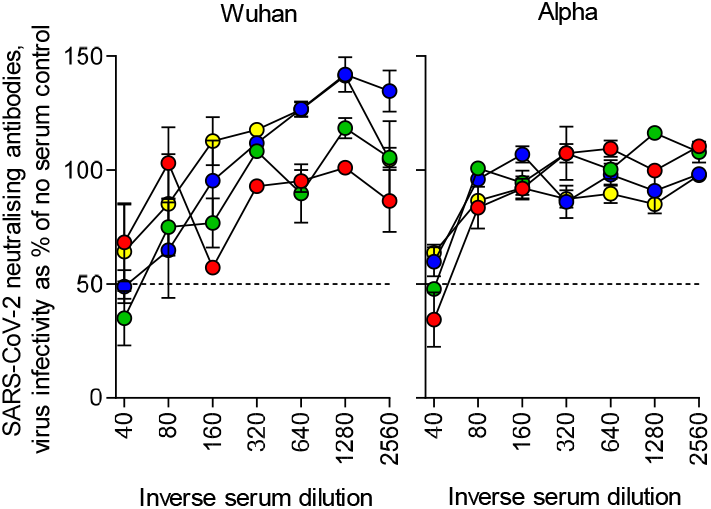
Prior HCoV-OC43 S immunity cross-reacts with SARS-CoV-2 S. Neutralisation curves of sera from mice after two doses of a HCoV-OC43 S DNA vaccine (n=4) of authentic SARS-CoV-2 Wuhan strain (left) or Alpha VOC (right), determined by a plaque reduction neutralisation test.

**Figure S2.**
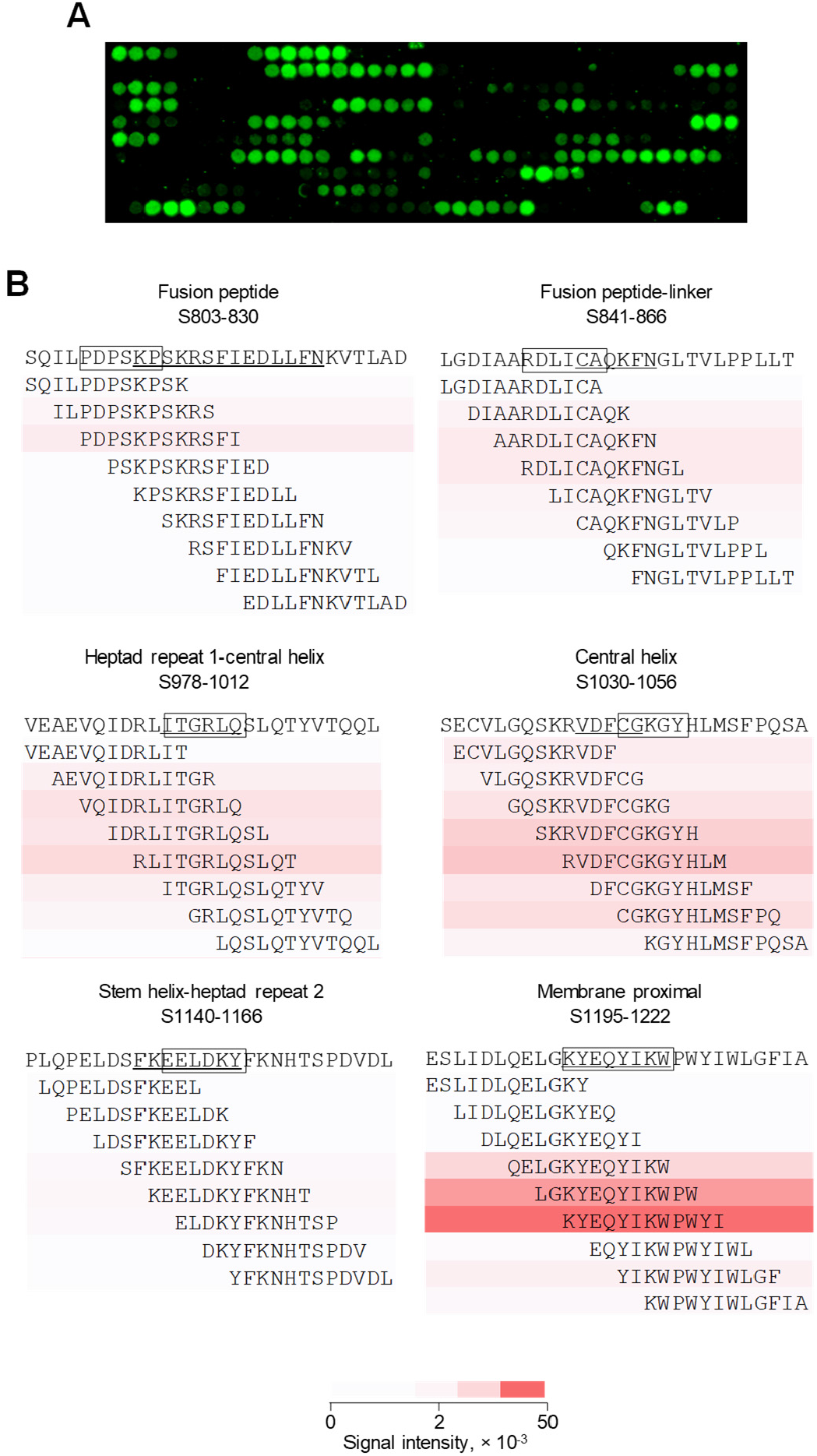
Mapping of cross-reactive epitopes in SARS-CoV-2 S using peptide arrays. **A**, Scanned image of a peptide array spanning the last 743 amino acids of SARS-CoV-2 S detected with sera from SARS-CoV-2 S2-immunised mice. The signal of the secondary antibody label (IRDye 800CW) is shown in green. The top left position in the array is the first peptide in the sequence (S_531-542_). The 12-mer peptides were arranged from left-to-right and top-to-bottom with an overlap of 10 amino acids, creating 367 spots. **B**, Heatmaps of signal intensity for the individual peptides spanning previously defined epitopes in the indicated S2 regions (underlined sequence). The core epitopes recognised by immune mouse sera in this array are shown in boxes.

**Figure S3.**
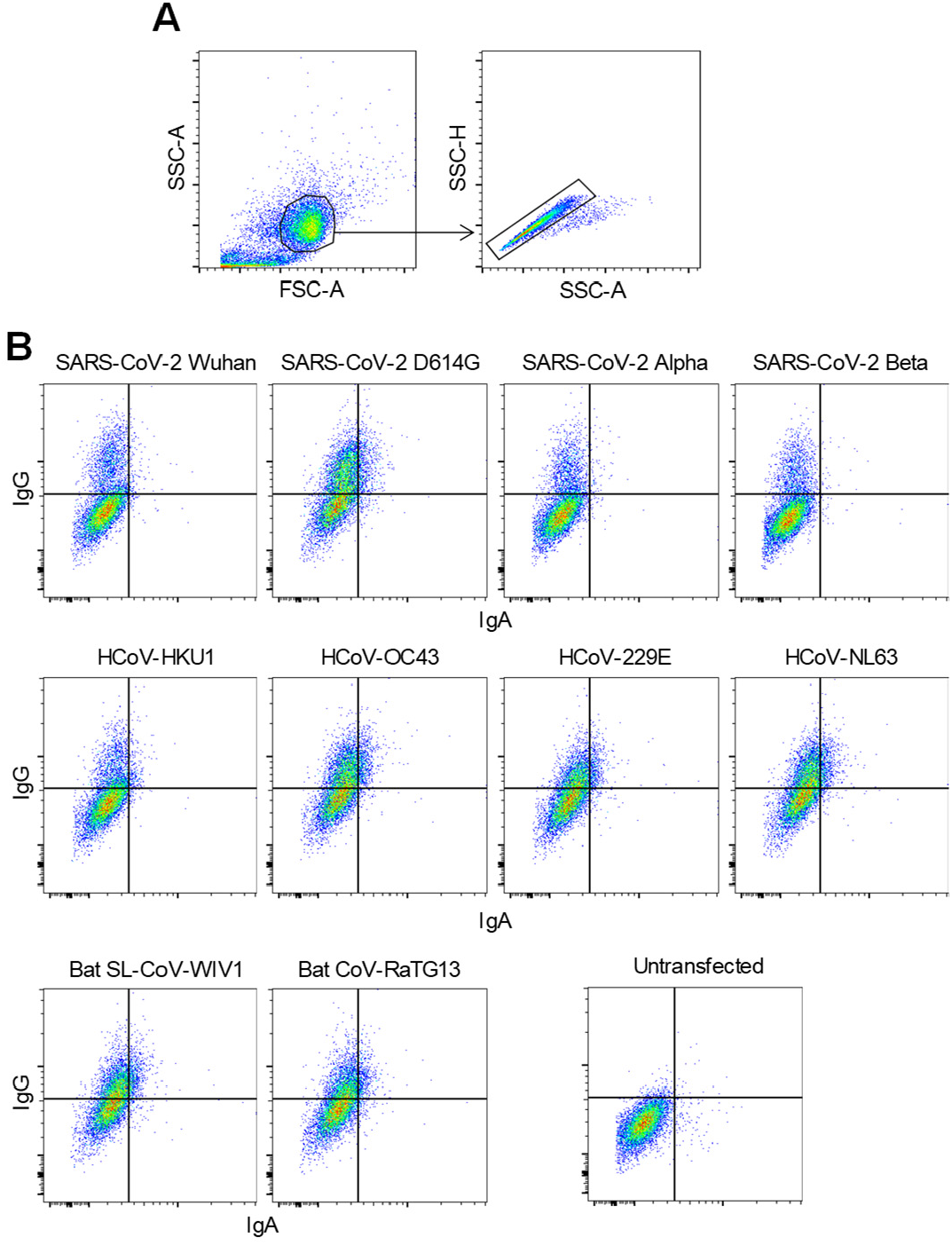
Gating of HEK293T cells expressing cell membrane-bound CoV S proteins. HEK293T cells were transfected with expression plasmids encoding the indicated CoV S proteins and were used for flow cytometric analysis two days later. **A**, Gating of HEK293T cells and of single cells in cell suspensions. **B**, Examples of staining with SARS-CoV-2 S2-immune mouse serum of HEK293T cells expressing each CoV S protein or negative control (untransfected) HEK293T cells.

**Figure S4.**
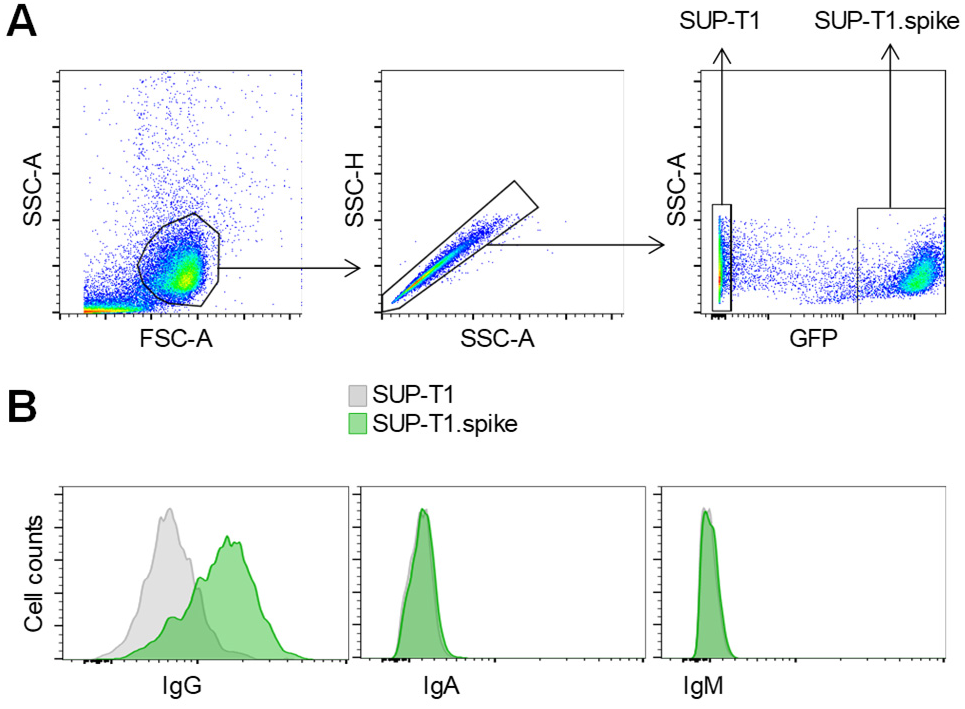
Gating of SUP-T1 cells stably expressing cell membrane-bound SARS-CoV-2 S. SUP-T1 cells, transduced with a retroviral vector encoding SARS-CoV-2 spike and GFP (SUP-T1.spike), were mixed with parental SUP-T1 cells in equal ratios and were used for flow cytometric analysis. **A**, Gating of SUP-T1 and SUP-T1.spike, according to GFP expression in the latter. **B**, Examples of staining with SARS-CoV-2 S2-immune mouse serum of these SUP-T1 cell mixtures.

**Figure S5.**
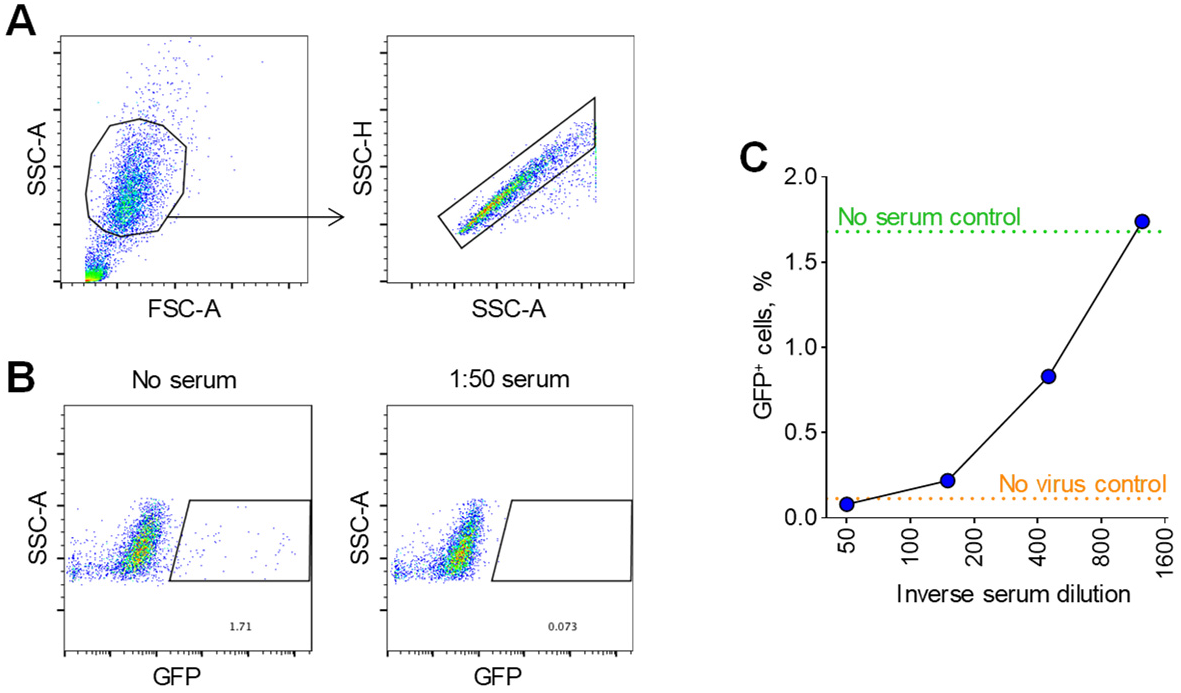
CoV S pseudotype virus neutralisation in Vero E6 cells. Vero E6 cells were transduced with CoV S pseudotypes encoding GFP. **A**, Gating of Vero E6 cells and of single cells in cell suspensions. **B**, Examples of GFP expression in Vero E6 cells transduced with a HCoV-OC43 S pseudotype in the absence (no serum) or the presence (1:50 serum) of SARS-CoV-2 S2-immune mouse serum. **C**, HCoV-OC43 S pseudotype neutralisation by the indicated dilutions of COVID-19 convalescent serum, used as a positive control. Dotted lines represent the % GFP^+^ Vero E6 cells obtained without serum (no serum control) or without transducing virus (no virus control).

